# Entrainment of theta, not alpha, oscillations is predictive of the brightness enhancement of a flickering stimulus

**DOI:** 10.1101/239061

**Authors:** Jennifer K. Bertrand, Nathan J. Wispinski, Kyle E. Mathewson, Craig S. Chapman

## Abstract

Frequency-dependent brightness enhancement, where a flickering light can appear twice as bright as an equiluminant constant light, has been reported to exist within the alpha (8 – 12 Hz) band. Could oscillatory neural activity be driving this perceptual effect? Here, in two experiments, human subjects reported which of two flickering stimuli were brighter. Strikingly, 4 Hz stimuli were reported as brighter more than 80% of the time when compared to all other tested frequencies, even though all stimuli were equiluminant and of equal temporal length. Electroencephalography recordings showed that inter-trial phase coherence (ITC) of theta (4 Hz) was: 1) Significantly greater than alpha, contralateral to the flickering stimulus; 2) Enhanced by the presence of a second ipsilateral 4 Hz flickering stimulus; and 3) Uniquely lateralized, unlike the alpha band. Importantly, on trials with two identical stimuli (i.e. 4 Hz vs 4 Hz), the brightness discrimination judgment could be predicted by the hemispheric balance in the amount of 4 Hz ITC. We speculate that the theta rhythm plays a distinct information transfer role, where its ability to share information between hemispheres via entrainment promotes a better processing of visual information to inform a discrimination decision.

## INTRODUCTION

When asked to judge the brightness of a stimulus, decisions regard the objective amount of light entering the eye (luminance) but are ultimately based on our subjective experience of this luminance (brightness). Despite our usual impression that subjective experience maps to objective reality, there can often be an enlightening disconnect between the two. As Von Helmholtz^1^ articulated, “…those cases in which external impressions evoke conceptions which are not in accordance with reality are particularly instructive for discovering the law of those means and processes by which normal perceptions originate”. A particularly strong example of this disconnect is the reported Brücke effect, where a stimulus flickering around 10 Hz requires half the luminance of a constant stimulus for the two stimuli to appear at equal subjective brightness^2^. This frequency-dependent brightness enhancement^2^ occurs over a range of slower frequencies (between 1 - 17 Hz).

Echoing Von Helmholtz^1^, the Brücke effect presents a compelling case where we can investigate the “…processes by which normal perceptions originate”. Here we probe the origins of brightness perception and tie the brightness-enhancement effect to changes in neural oscillations measured using electroencephalography (EEG). Bartley^2^ found the greatest difference between luminance and brightness when stimuli flickered near 10 Hz, in the alpha frequency range. Described as an inhibitory internal sampling rhythm^3-5^, alpha oscillations have been shown to impact a diverse array of visual percepts. For example, visual detection can be predicted by the prestimulus power and phase of endogenous alpha rhythms^6^, disrupted or reset alpha rhythms^7^, and entrained alpha rhythms^8-10^. A recent review of EEG studies^11^ suggests that these alpha dependent changes to visual perception are largely phasic, where one phase of the oscillation (e.g., a peak) favours one outcome (e.g., detecting a target), and the opposite phase (e.g., a trough) favours the other outcome (e.g., failing to detect a target).

In 1966, Kohn and Salisbury^12^ replicated the Brücke effect on brightness reports, but concluded there was no connection to their EEG measures. However, they used only 2 participants and 2 electrodes with an analysis of oscillatory content that included simply measures of power (amplitude) and not phase (timing) - limitations that likely precluded them from observing the effects we report here. Interestingly, while Kohn and Salisbury^12^ discuss their findings as showing maximal brightness enhancement for ∼10 Hz flickering stimuli, a closer examination of their results (p. 464; fig. 3) suggests that they saw the largest effects with stimuli flickering at a slower rate close to 5 Hz. Perhaps changes to oscillations in the theta rhythm (around 3-8 Hz) may instead be responsible for brightness enhancement.

Like the alpha band, theta phase and power, both pre- and post-stimulus, have proven to play a role in various visual percepts, from attentional search^13^, to contrast gain^14^, and even to subjective colour preference^15^. Further, and of particular relevance to the current study, recent work from Han & VanRullen^16,17^ shows that the phase of theta oscillations plays a role in the apparent brightness enhancement of a meaningful image as opposed to a random line image. They suggest the appearance of the fully connected and interpretable line object as brighter than the random line image is due to theta oscillations carrying top-down signals from higher areas when making luminance judgments. This interpretation is supported by a study of contrast judgments, which were most difficult when masked by a frequency just faster than theta^18^. According to the authors, and in line with the results of Han & VanRullen^16,17^, a mask “slightly faster than the perceptual cycle” governed by theta optimally disrupted judgments in this experiment^18^.

Therefore, we designed the current study to answer two main questions: First, in replicating the Brücke effect by testing frequencies within the original reported range^2^ of brightness enhancement (0 – 17 Hz), would we find the largest effect for stimuli flickering in the theta (4-5 Hz) or alpha (9-10 Hz) range? Second, regardless of what frequency induced the largest brightness enhancement, would this be reflected in the phasic response of the EEG oscillations of that frequency? To answer these questions, we ran two experiments. First, we conducted a behavioural experiment designed to replicate the Brücke effect (one flickering stimulus compared to one constant stimulus and also extend it to two flickering stimuli, with all flickering stimuli appearing in the range of the originally reported enchancement^2^). Second, we used the results of the first experiment to refine our frequency range to those providing the greatest brightness enhancement and examined phasic, oscillatory effects at the frequencies of the flickering stimuli by measuring their inter-trial phase coherence (ITC). In brief, in both experiments we found that a 4.4 Hz flickering stimulus elicited the greatest apparent brightness enhancement – chosen as brighter over the other frequency stimuli more than 80% of the time. In the EEG study, we found that this 4.4 Hz brightness enhancement was paired with the greatest amount of phase locking, measured with ITC.

## RESULTS

### Choice behaviour

#### Flicker at theta frequency produces the greatest behavioural brightness-enhancement effect

In both Experiment 1 (E1) and Experiment 2 (E2) participants were asked to judge which of two stimuli on the screen was brighter (or darker, question counterbalanced across participants). We presented a range of frequencies equally spaced across the originally reported range of enhancement^2^ in E1 (0, 4.4, 9.2, 13.3, 17.1 Hz) and a restricted subset in E2 (4.4, 9.2, 13.3 Hz) in all possible pairs (Figure 1) and recorded the side and frequency of the stimulus participants selected as brighter on each trial. The results for E1 and E2 can be seen in Figure 2 (panels A & C, and B & D, respectively).

**Figure 1:**
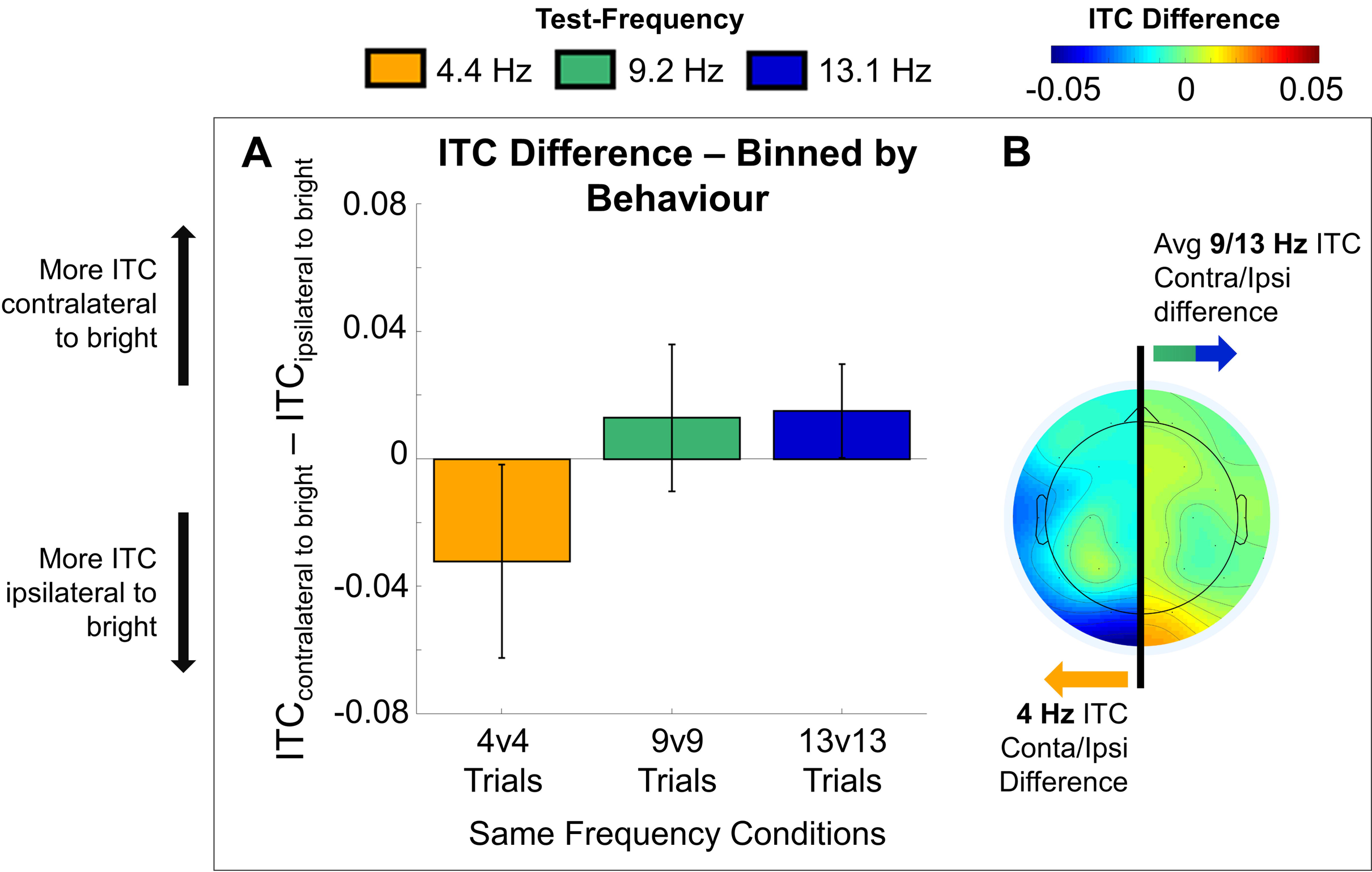
The timing of a single trial of E1 and E2, with stimuli dimensions. Initial fixation was a variable length, randomly jittered (between 750 and 1250 ms in E1, and between 1250 and 1750 ms in E2). Flashing stimuli, the entrainment and response portion, occurred for 4000 ms in both E1 and E2, followed by an inter-trial interval (ITI) of 500 ms for E1 or 560 ms for E2.

**Figure 2:**
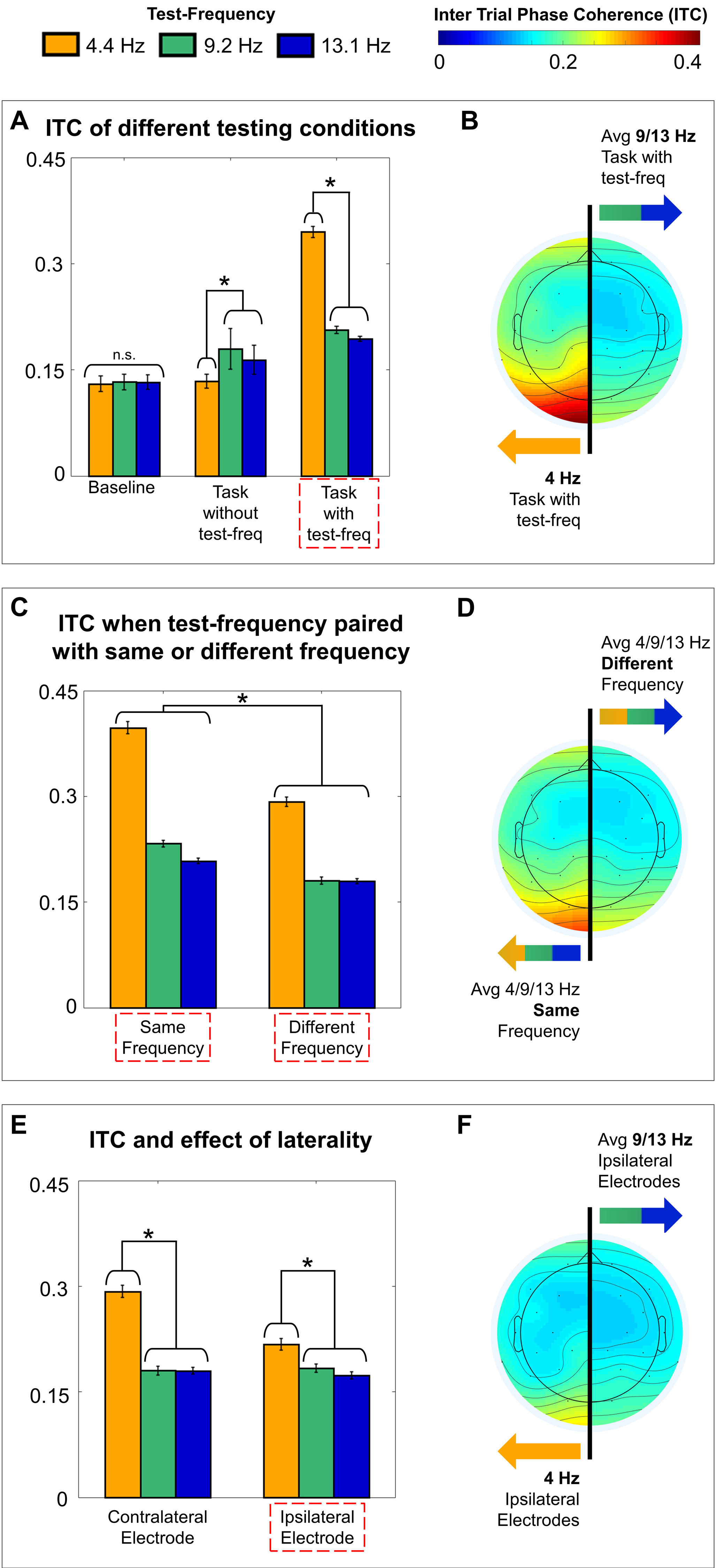
**A**E1 choice proportion of choosing the left stimulus frequency as brighter over the right stimulus frequency (as a subtraction) **B** E2 choice proportion of choosing the left stimulus frequency as brighter over the right stimulus frequency (as a subtraction). For A and B, white denotes 50% choice proportion, pink colours represents a greater choice proportion of the left stimulus frequency, and purple colours denotes a greater choice proportion of the right stimulus frequency. **C** E1 choice proportion of choosing the test-frequency (on x axis) over all frequencies, separated by the side the test-frequency appeared on (black = left, grey = right). **D** E2 choice proportion of choosing the test-frequency over all frequencies, separated by side of space, following the same conventions as plot B. For C and D, error bars represent the average of individual standard errors.

Critically, when paired with a stimulus of any other frequency, 4.4 Hz stimuli were chosen as brighter more than 80% of the time. This was confirmed via a two-factor repeated-measures (RM) analysis of variance (ANOVA) comparing left versus right frequencies (E1: 5 frequencies x 5 frequencies; E2: 3 frequencies x 3 frequencies). In both experiments, we found a main effect for both Left Frequency (E1: *F*_(0.41,11.37)_ = 26.0; *p* = 3.1e-07; ε= .10; E2: *F*_(0.65,14.26)_ = 59.5; *p* = 3.2e-11; ε= .32) and Right Frequency (E1: *F*_(0.41,11.37)_ = 30.3; *p* = 8.7e-09; ε= .10; E2: *F*_(0.65,14.26)_ = 49.6; *p* = 4.4e-10; ε= .32) and no significant interaction (see Table 1 for E1 and E2 choice behaviour pairwise comparisons). The lack of an interaction indicates there was no systematic side of space bias. Less dramatically, in E1, 9.2 Hz stimuli were chosen as brighter significantly more than 13.3 and 17.1 Hz, and 0 Hz stimuli were chosen as brighter more than 17.1 Hz stimuli. In E2, we noticed a similar trend for 9.2 Hz to be selected as brighter than 13 Hz, but this did not reach significance. In E1, the 0 and 9.2 Hz stimuli, and the 13.3 and 17.1 Hz stimuli were not selected at significantly different rates. Finally, in conducting these follow-ups in E1 we found a very small side of space bias that existed between 0 and 13.3 Hz, where only a 0 Hz stimulus on the right was chosen as brighter more often, while on the left, though 0 Hz was selected as brighter more often, this was not significant.

**Table 1:**
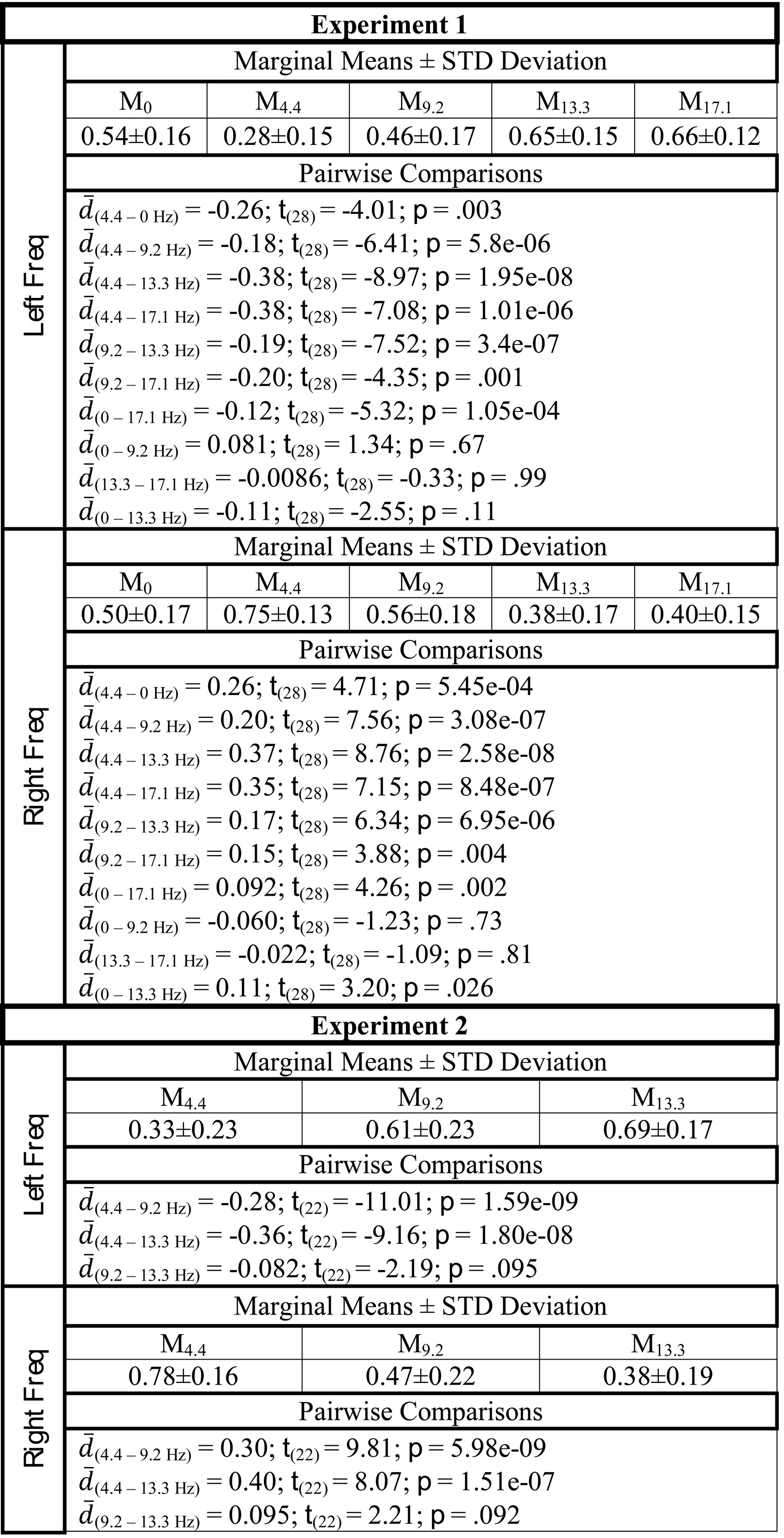
Marginal (simple main effect) means, standard deviations and significant pairwise comparisons for choice behaviour in E1 and E2. Pairwise comparisons are reported with mean differences (specified by subscript), t statistics, and the corresponding *p* value. Reported are all significant pairwise comparisons as well as pairwise comparisons that did not achieve significance but were discussed in the text.

### Neural oscillations

To test for phase-related effects that correlated with the observed brightness enhancement, in E2 we calculated the across-trial phase coherence, or ITC, in each of three frequency bands (4.0-5.0, 8.5-9.5, and 13.0-14.0; all units Hz) for the time window 1-2 s after the flickering stimuli first appeared on the screen (except for a baseline period, see next section). All statistically analyzed ITC values were extracted from occipital electrodes O1 and O2, though some of the reported effects also manifest dorsally along the midline (e.g., see topographies in Figure 3). We analyzed these ITC values to answer four specific questions: First, would the pattern of ITC follow the pattern of choice behaviour such that ITC contralateral to 4.4 Hz flickering stimuli is larger than ITC contralateral to the 9.2 or 13.3 Hz stimuli? Second, in an effort to further explore the question of how the brain processes two flickering stimuli at the same time, would ITC values for two stimuli of the same frequency be larger than ITC values contralateral to a different-frequency stimuli pair? Third, would there be any evidence for intra-hemispheric effects such that the ITC ipsilateral to a flickering stimulus would be modulated by its frequency? Fourth, if we examine only trials with identical stimuli (4.4 vs 4.4 Hz, 9.2 vs 9.2 Hz and 13.3 vs 13.3 Hz) sorted by participant response, would there be frequency-specific ITC markers that predict which stimuli was perceived as brighter?

**Figure 3:**
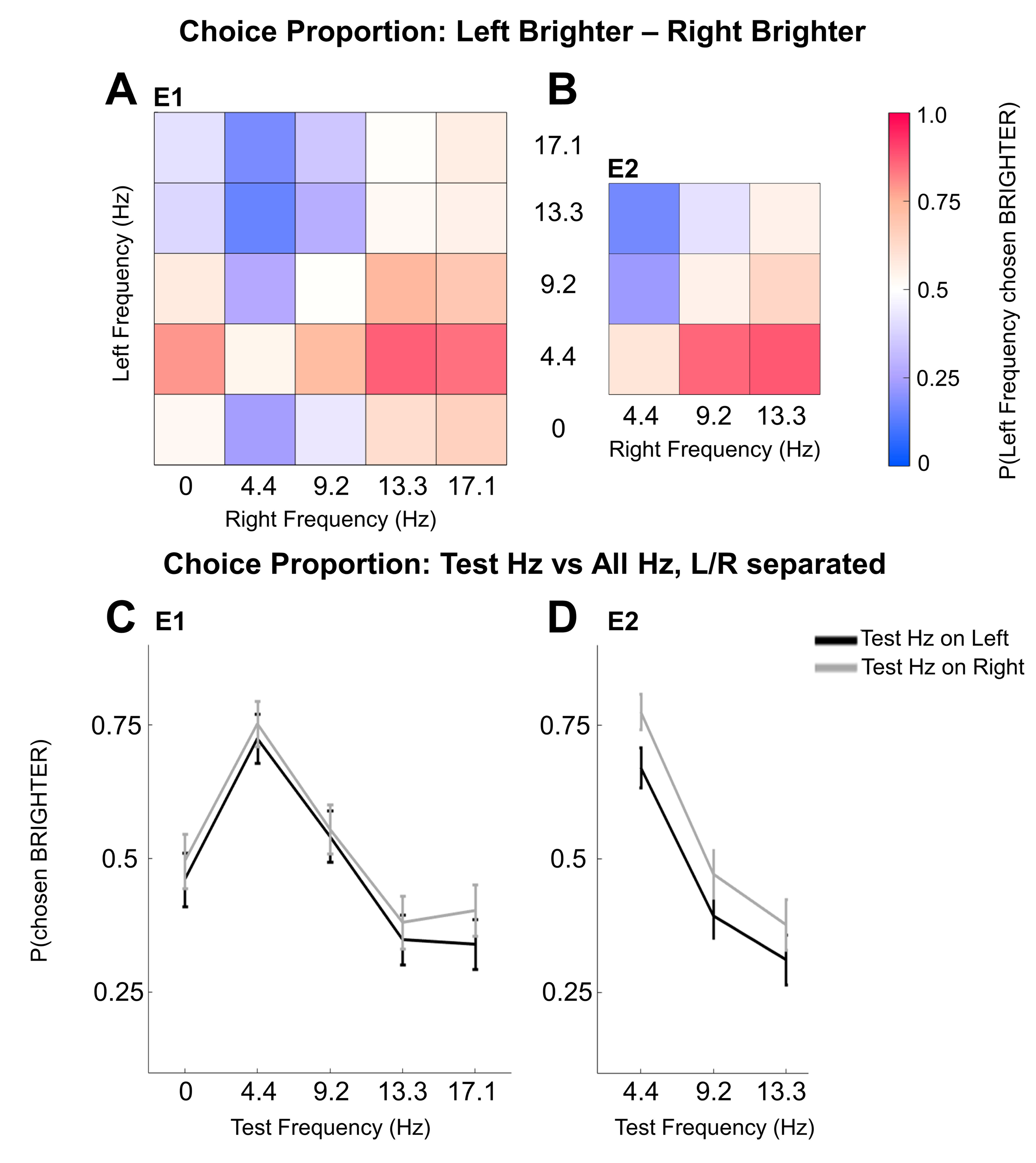
ITC results. Bar plots (A,C,E) represent ITC data of the test frequency (see legend) from only occipital electrodes (O1/O2). Error bars represent averaged individual standard error and * indicates where pairwise comparisons were significant. Topoplots (B,D,F) show the ITC of critical conditions (red boxes on bar plot axes), one on each side of the brain. **A** ITC of different testing conditions: Baseline (no stimuli on the screen); Task without test-frequency (Stimuli but not the test-frequency on the screen) Task with test-frequency (Stimuli including the test-frequency on the screen, measured contralateral to the test-frequency). **B** The topoplot of the Task with test-frequency result, showing 4 Hz band ITC on the left, and an average of 9 and 13 Hz bands ITC on the right. **C** ITC when the test-frequency was paired with a stimulus of the same frequency or a different frequency. **D** The topoplot showing on the left, the average ITC of all test frequencies when paired with a stimulus of the same frequency, and on the right the average ITC of all test frequencies when paired with a stimulus of a different frequency. **E** ITC from electrodes contralateral and ipsilateral to the test frequency stimulus. **F** The topoplot of ipsilateral electrode responses showing amount of 4 Hz band ITC on the left, and average amount of 9 and 13 bands Hz ITC on the right.

#### ITC contralateral to 4.4 Hz flickering stimuli is largest

We calculated ITC values in the three frequency bands of interest (referred to as test-frequency) across three different conditions: 1) baseline (before stimuli onset, ITC was averaged across hemispheres since no stimuli were on the screen), 2) while stimuli were on the screen but the test-frequency was not (ITC was averaged across hemispheres since the test-frequency was not on the screen), and 3) while stimuli were on the screen including the test-frequency (ITC was measured contralateral to the test-frequency). We ran a 2 factor 3 × 3 (Frequency x Condition) RMANOVA, with the results depicted in Figure 3A-B. Both main effects of Condition, (*F*_(0.43,9.40)_ = 58.6; *p* = 1.9e-08; ε = .21) and Frequency, (*F*_(0.43,9.40)_ = 15.1; *p* = 1.5e-04; ε = .21) were significant, as was their interaction, (*F*_(0.85,18.80)_ = 30.3; *p* = 1.9e-06; ε = .21). Follow up simple main effect RMANOVAs were carried out across Frequency at each level of Condition. There was no significant effect of Frequency in the baseline Condition (*F*_(1.95,42.97)_ = 0.46; *p* = .63; ε=.98) indicating that prior to stimuli, there were no frequency-specific differences in ITC. For the condition where there were stimuli on the screen, but not the test-frequency, there was an effect of Frequency (*F*_(1.95, 42.91)_ = 18.91; *p* = 9.7e-07; ε= .98) with frequency-specific increases for 9.2 compared to 4.4 Hz stimuli 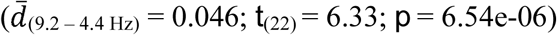 and 13.3 compared to 4.4 Hz stimuli 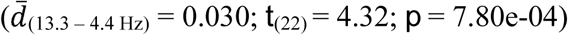. Finally, for the critical condition of stimuli within the test-frequency being on the screen, we found a significant effect of Frequency (*F*_(1.11,24.38)_ = 19.81; *p* = 1.5e-05; ε = .55), and multiple comparisons revealed that 4.4 Hz had significantly larger ITC values compared to 9.2 (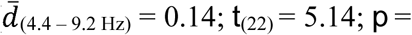 1.07e-04) and 13.3 Hz 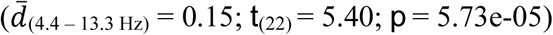. These results indicate: First that at baseline, no frequency naturally has greater phase locking; Second that the task itself appears to elicit an increase of 9.2 and 13.3 Hz ITC (possibly task related attention – a point we return to in the discussion); and Third that 4.4 Hz stimuli show increased contralateral ITC values in the respective test-frequency compared to the other two frequency stimuli. This then provides evidence that a neural correlate of the brightness enhancement observed in the Brücke effect is the degree to which a visual stimulus induces theta-band phase locking in occipital electrodes.

#### Two same-frequency stimuli elicit larger ITC than two different-frequency stimuli

In the critical condition described above, with stimuli on the screen within the test-frequency, the data collapses across trials in which the test-frequency appears with itself (e.g. 4.4 vs 4.4 Hz) and trials in which the test-frequency is paired with a different frequency (e.g. 4.4 vs 9.2 Hz). To explore the effects of having two of the same-frequency stimuli on the screen, a 2 factor 3 × 2 (Frequency x Stimulus Type) RMANOVA compared the ITC contralateral to the test-frequency when the stimuli were the same (Same-frequency) or different (Different-frequency). The results of this analysis are shown in Figure 3C-D. Both Stimulus Type, (*F*_(0.34,7.59)_ = 32.1; *p* = 1.1e-05 ε; = .34), and Frequency, (*F*_(0.69, 15.18)_ = 27.0; *p* = 1.5e-05; ε = .34) showed significant main effects, as did their interaction, (*F*_(0.69, 15.18)_ = 11.0; *p* = 4.6e-04; ε = .34). For all levels of Frequency, the effect of Stimulus Type was significant (4 Hz: *F*_(1,22)_ = 41.29; *p* = 1.8e-06; 9 Hz: *F*_(1,22)_ = 13.03; *p* = .001; 13 Hz: *F*_(1,22)_ = 5.10; *p* = .03) with ITC showing larger values for Same-frequency than Different-frequency 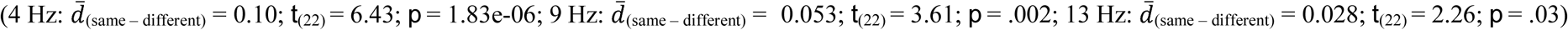. This shows that two lateralized stimuli do affect each other, and that two stimuli of the same-frequency will promote more phase locking, despite being presented in opposite visual hemi-fields.

#### Ipsilateral increases to ITC seen only for 4.4 Hz stimuli

Again, we further narrowed our ITC analysis and examined only the “different-frequency” trials from the second ITC analysis. To explore how the whole brain response might be implicated, we compared the electrode responses ipsilateral versus contralateral to the test-frequency and ran a 3 × 2 (Frequency x Side of Electrode [Contralateral/Ipsilateral]), 2 factor RMANOVA. We found main effects for Frequency, (*F*_(0.69,15.11)_ = 20.1; *p* = 5.6e-05; ε = .34) and Side of Electrode, (*F*_(0.34_, _7.55)_ = 16.8; *p* = 4.7e-04; ε = .34) as well as a significant interaction, (*F*_(0.69,15.11)_ = 7.3; *p* =01; ε = .34). The results of this test are shown in Figure 3E-F. Follow up simple main effect RMANOVAs were carried out across Frequency and found significant Frequency differences for both the Contralateral Electrode (*F*_(1.09,23.92)_ = 17.35; *p* = 2.6e-04; ε = .54) and the Ipsilateral Electrode (*F*_(1.55,34.09)_ = 7.20; *p* = .005; ε = .77). Post-hoc comparisons demonstrated that for both Contralateral and Ipsilateral, these differences were driven by higher 4.4 Hz ITC values compared to 9.2 Hz (Contralateral: 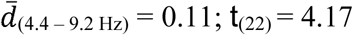; *p* = .001; Ipsilateral: 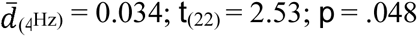) and 13.3 Hz (Contralateral: 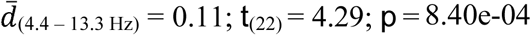; Ipsilateral: 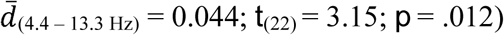. This analysis confirms the finding that 4.4 Hz stimuli induce more phase locking than 9.2 and 13.3 Hz stimuli in contralateral electrodes and further suggests that a 4.4 Hz stimuli is capable of inducing ipsilateral phase locking. To now, these results suggest that there are two possible mechanisms at play for brightness enhancement. While it appears definitive that theta-band ITC is correlated with brightness enhancement, it remains unclear whether this effect is driven more by contralateral phase locking, or more by the ability for the phase locking to be shared across hemispheres and thus to also be represented in the ipsilateral electrode.

#### Brightness discrimination behaviour predicted by frequency specific ITC changes

For this analysis, we ask, when the stimuli presented are identical and the bias to choose one target over the other is entirely internal, is this bias reflected in our measures of brain activity? Thus for this fourth and final test of ITC we examined exclusively the “same-frequency” trials but sorted the trials by the choice response such that we averaged across electrodes that were contralateral or ipsilateral to the stimuli selected as brighter. For this specific analysis, since we didn’t know at what time internal biases might be generated, we expanded our ITC window, and calculated over the range of possible response times (100-3212 ms).

We tested whether the brightness enhancement induced by theta-band phase locking was due more to local processing, (e.g. electrodes contralateral to the “brighter” stimulus show more ITC) or due more to global processing (e.g. electrodes ipsilateral to the “brighter” stimulus show more ITC). This was most concisely represented as a contralateral-to-brighter minus ipsilateral-to-brighter ITC difference. Moreover, we included all same-frequency trials to also test whether there were any hemispheric ITC effects in the other frequency bands (e.g. 9.2 and 13.3 Hz) that were predictive of choice behaviour (see Figure 4A).

**Figure 4:**
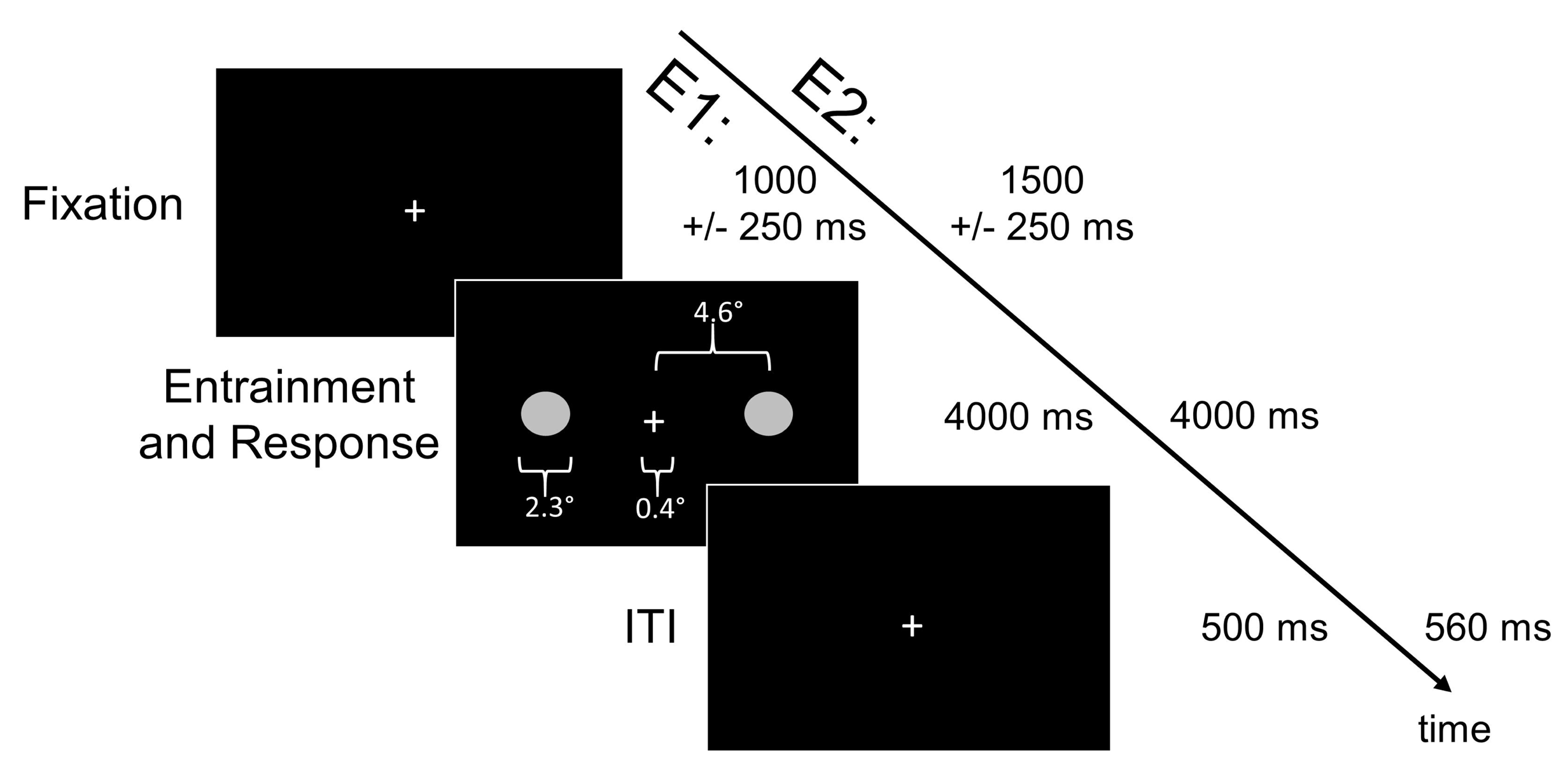
Brightness discrimination on same-frequency trials predicted by frequency-specific ITC changes. **A** Bar plot representing the difference in behaviour-binned ITC between the occipital electrode contralateral to the stimulus chosen as brighter and the occipital electrode ipsilateral to the stimulus chosen as brighter. A negative value denotes more ITC ipsilateral to stimuli judged as brighter and a positive value denotes more ITC contralateral to a stimuli judged as brighter. Error bars represent the standard error of the mean difference for each test-frequency. **B** The topoplot corresponding to differences depicted in A, showing the 4 Hz band ITC difference on the left, and the average of the 9 and 13 bands Hz ITC difference on the right.

We ran a one-factor, RMANOVA on the contralateral-ipsilateral to brighter ITC difference and found a significant main effect of Frequency, (*F*_(1.60,35.27)_ = 7.4; *p* = .004; ε = .80). Multiple comparisons showed that this effect was driven by significant differences between both 4.4 and 9.2 Hz 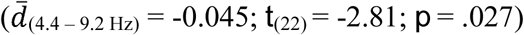 and 4.4 and 13.3 Hz 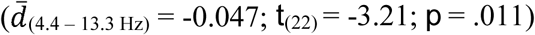. In addition to these post-hoc comparisons, we separately analyzed the direction of this difference by using one sample t-tests to compare each frequency specific effect against zero (error bars in Figure 4A). Interestingly, the 4.4 Hz effect is significantly negative (*t*_(22)_ = −2.20; *p* = .039) while the 9.2 Hz and 13.3 Hz are numerically positive, with the 13.3 Hz effect achieving significance (*t*_(22)_ = 2.13; *p* = .045). This pattern suggests that in the theta band, it is the global sharing of information across hemispheres that is most important to perceiving a stimuli as brighter (ipsi > contra) while in the alpha band (9-13 Hz) it is the local response that is most important to perceiving a stimuli as brighter (contra > ipsi).

## DISCUSSION

This study was motivated by the pronounced dissociation between luminance and brightness judgments induced by flickering stimuli^19-25^. Here we aimed to answer two questions: 1) Would we find the largest brightness enhancement for stimuli flickering in the theta (4-5 Hz) or alpha (9-10 Hz) range? And 2) Would this brightness enhancement be reflected in the phasic response of the EEG oscillations of that frequency? First, in two experiments designed to test the originally reported range of frequencies (0-17 Hz) known to elicit brightness enchancement^2^, we found that a stimulus flickering at 4.4 Hz was perceived as brighter more than 80% of the time when compared to any other flickering or constant (0 Hz) stimulus within the 0–17.1 Hz range (Figure 2). This was true even though all stimuli were equiluminant and presented for the same length of time. Second, in E2 we recorded EEG and focussed on the measure of inter-trial phase coherence, or ITC. Most importantly, ITC in the 4.4 Hz band was highest for electrodes contralateral to 4.4 Hz stimuli, exactly matching the large behavioural effects we report. Thus, we have clear evidence that a neural correlate of frequency-induced brightness enhancement is theta-band phase locking.

We were able to confirm that this ITC effect was not due to a general increase in 4.4 Hz band ITC (e.g. during baseline) nor was it due to a task (but not frequency) related increase in 4.4 Hz band ITC (though, this increase did occur for 9.2 and 13.3 Hz stimuli, a point we return to later). Further, we showed that two same-frequency stimuli induced more phase locking than two different-frequency stimuli (especially for two 4.4 Hz stimuli) and that 4.4 Hz stimuli are more successful at inducing both contralateral *and* ipsilateral phase locking (Figure 3). This ability for a 4.4 Hz signal to propagate across hemispheres may be a key factor in brightness enhancement. This hypothesis was directly tested by examining the ITC response on same-frequency trials (e.g. visual stimulation in both hemi-fields was identical) in electrodes contralateral to stimuli chosen as brighter versus those ipsilateral to stimuli chosen as brighter. Here, for 4.4 Hz stimuli the ipsilateral-to-brighter electrode showed more ITC while for 9.2 and 13.3 Hz stimuli the contralateral-to-brighter electrode showed more ITC (Figure 4).

An outstanding question arising from our results is why we did not see brightness enhancement (nor any neural correlate of it) in the alpha band. This is surprising not only because the original report found maximal enhancement around 10 Hz^2^, but it also seems to stand in contrast to the rich literature on the effects of alpha oscillations on perception (for a review, see^26-27^). First, whereas we used a fast-refresh LCD monitor to display our flickering stimuli, Bartley used a rotating disk to periodically block a constantly lit light bulb. This could have changed the (what he referred to as) “light-to-dark ratio” (LDR) – essentially the duty cycle of the flickering stimulus. On this point, however, Bartley did probe a variety of different LDRs and reported an ∼ 10 Hz peak for all ratios. Second, it is not clear that Bartley tested anyone other than himself (other than one comment that the results from a 12-year-old “compared favourably with the author’s”), and also, it is unclear how many trials contributed to the results. Finally, while we get a good deal of consistency across our participants, there are known individual differences in other naturally occurring brain rhythms (e.g. variability of alpha band^28^) so it is possible that Bartley’s report was idiosyncratic to him. It is also important to recall that the 1966 replication of the Brücke effect with limited EEG by Kohn and Salisbury^12^ actually had some of the largest brightness enhancement in the 5 Hz, and not the 10 Hz, range.

Here we propose a speculative, but more intriguing reason for the discrepancy between previous reports of alpha-band brightness enhancement and the current report of theta-band brightness enhancement. As a caveat to this speculation, we do not assume causality in the relationships between specific oscillatory activity and associated perceptual phenomena. While we understand EEG represents a reading of systems or ensembles of neurons, it must only be interpreted as a reflection of the underlying processes, and we cannot yet speak to the neural basis of these processes (or suggest processes are categorically exclusive of other processes) beyond documenting phenomena associated with them. Further, we understand that many different frequencies of oscillations have been found to correlate with many different cognitive and perceptual functions, and the scale of these oscillatory frequencies may differ greatly between EEG findings and single cell recordings. That being said, we believe it may be the nature of the task, and the judgement required, that separates whether the phase and power of alpha or theta rhythms will have the greatest impact on performance. Effects of neural oscillations on perception fall into one of two clusters, either near 10 Hz (referred to as alpha) or near 7 Hz (referred to as theta)^11^. Alpha effects are related to sensory sampling, and theta effects with higher order “attentional” sampling^11^. Here we offer a refinement of that idea – if the nature of the task demands only the ability to report *if* a sensory event occurred, then it is likely to be governed by the alpha rhythm. However, if a task requires the identification of *what* a stimulus is, the behaviour will be most impacted by theta rhythms. For example, in detection tasks^6^ or single item discrimination tasks^29^, success is based almost exclusively on whether or not you acquired the low-level, bottom-up visual information from the outside world. Thus, it makes sense that the sensory-sampling rhythm, alpha, is most relevant. Of note, the constant comparison stimuli in the original report of the Brücke effect^2^ might therefore fall in this domain. However, in higher-order tasks, like the discrimination task used in the current experiment, where it is necessary to store and/or compare information, we argue that the theta rhythm is most relevant. Under this framework, alpha oscillations are responsible for periodically extracting sensory information from the world while theta oscillations function as a transfer rhythm, or carrier wave^30^ to share that information between brain areas either for additional processing or comparison. With the understanding of oscillations as cycles that alternate between periods of excitability and inhibition^31^, where the cycles can coordinate and synchronize neural activity within and between brain areas, alpha oscillations facilitate sensory information sampling within more local areas, and theta oscillations display a periodicity of excitation that provides long-range communication between brain areas. With respect to our task, and why 4.4 Hz therefore looks brighter, we believe that theta periodicity in the stimuli creates the most optimal information transfer by way of aligning the naturally-occurring endogenous theta rhythm and its long-range connections to a temporally strict and precise schedule. This precision allows for optimal information transfer resulting in a bias to see that stimulus as brighter when compared to a second target. In contrast, for stimuli oscillating close to the alpha rhythm (e.g. 9.2 or 13.3 Hz), while they may have an advantage in being detected, their periodicity interferes with the theta rhythm, putatively resetting its phase before optimal throughput has occurred. Thus, with a disrupted flow of information, these stimuli are not perceived as bright as their 4.4 Hz counterpart.

Our analysis of the same-frequency trials binned by behavioural response supports this framework. Specifically, if periodic stimuli presented in the alpha range optimize sensory sampling, then one would predict that brighter stimuli would be those that were the most successfully entrained at the site of response. Consistent with this idea, the 9.2 vs 9.2 Hz and 13.3 vs 13.3 Hz trials in E2 showed elevated stimulus related ITC (e.g. 9.2 Hz band and 13.3 Hz band) in electrodes contralateral to the stimulus chosen as brighter, even though the two stimuli were identical. Conversely, if periodic stimuli presented in the theta range optimize information transfer, then one would predict that brighter stimuli would be those that were most successfully entraining a whole brain response. The 4.4 vs 4.4 Hz trials showed elevated 4.4 Hz band ITC in electrodes ipsilateral to the stimulus selected as brighter. Importantly, this final finding does not take away from the general finding that theta-band entrainment leads to brightness enhancement (e.g. the pattern of behaviour on different-stimuli trials), but rather, suggests that the theta-band dominance in our brightness enhancement task may arise because of its ability to entrain optimal information flow across hemispheres.

This framework of complementary but distinct roles for alpha and theta oscillations can also account for theta’s information sharing role in a variety of higher order processes, from attention^13^, to movement^32^ and its planning^33^, and to memory^28^. Further, evidence exists for theta’s facilitation of a brightness enhancement of meaningful stimuli, where no effect is seen for random line images, suggesting theta may also provide feedback from higher-order areas^16,17^. In contrast, alpha as a sensory sampler could also in part explain why we see task-related phase locking in the alpha (9.2 and 13.3 Hz bands) range in E2 (and has been previously reported as a hallmark of good visual discrimination performance^34^). That is, in a task where the arrival of visual information is predictable across trials, it makes sense for the brain to prepare to be maximally sensitive, and thus, phase lock the sensory sampling rhythms.

Importantly, there is an alternative explanation for theta-band effects as simply reflecting alpha sampling when the task requires more than one object to be sampled (as in our task). In this model, there is no distinct role for theta rhythms. Rather, alpha oscillations only acquire a sample from one “object” at a time, so, with two objects it optimally samples each one at half of alpha, or close to our 4.4 Hz. From our study, it is impossible to disentangle whether our finding in 4.4 Hz is truly the result of an oscillation in the 4.4 Hz band, or instead follows a more recently proposed model of a halving of the alpha rhythm when two stimuli are present^35-37^. However, while this could be a possible account of our results, it doesn’t appear to explain the variety of tasks listed above in which sensory sampling is required from only one^32^ or more than two^13^ locations, nor does it offer as intuitive an explanation for the finding that the electrodes ipsilateral to stimuli selected as brighter on 4.4 vs 4.4 Hz trials are those showing the highest phase locking. Future work could test these competing hypotheses (alpha / 2 vs. theta) as well as pursue other interesting lines of questioning (e.g. the effects of monocular viewing, and eye dominance^38^, as well as exploring even lower frequencies for enhancement).

Bartley provided one of the first documented reports of the Brücke effect – the powerful illusion where flickering light sources (in the 0-17 Hz range) appear brighter than constant light sources^2^. In that publication, Bartley also predicted that “the brain mechanism whose rhythmic activity determines the rate of cortical responses produces the Brücke effect”^2^. Impressively almost 80 years later, this appears to be true. While the entrainment of an alpha rhythm (9.2 Hz in this case) was the hypothesized source of the generation of a perception different from sensation, it now appears that it is the entrainment of the theta rhythm, our proposed information sharing rhythm, that provides this brightness enhancement. The consistency of phase angles, or the strength of phase-locking, in the theta rhythm goes hand in hand with brightness enhancement. Our study adds another percept to the list of behaviours correlated with theta activity, compounding the complexity of the discussion, and highlighting the work still needed for a complete understanding of this mysterious and pervasive rhythm.

## MATERIALS AND METHODS

### Subjects

A total of 30 individuals (11 males, mean age 27.2 +/− 8.8 years) participated in E1, with one participant excluded after determining their left-handedness post-experiment setup, resulting in 29 behavioural data sets for analysis. E2 involved a total of 23 individuals (7 males, mean age 19.1 +/− 1.4 years), with all participants not having participated in E1. All participants self-reported that they were right-handed and had normal or corrected-to-normal vision. No participant had prior knowledge about the experiment and its objectives. All experimental proceedings were approved by the University of Alberta’s Research Ethics Office (Pro00059044) and were performed in accordance with relevant guidelines and regulations. All participants gave informed consent prior to participating and were either paid for their time at a rate of $10 per hour or rewarded with credit toward their introductory psychology course.

### Apparatus

Participants sat at a table 50 centimeters (cm) away from a ViewPixx 120-Hz refresh rate monitor and a keyboard in a dimly lit room. The study was designed and carried out with a computer running Windows 7 and the Psychophysics^39^ toolbox in Matlab. The stimuli were two circles, one left and one right of the central fixation cross, and were presented binocularly in the full visual field. The circles were a diameter of 2.3°, with 4.6° between the centre of each circle and the centre of the fixation cross (width of 0.4°), as illustrated in Figure 1. The stimuli would flicker between black, the colour of the background, and grey, with a pixel value of [128 128 128]. Stimuli that would “flicker” at a frequency of 0 Hz were presented as a grey circle that remained on the screen for the entire stimulus duration.

### Tasks/Procedure

Each trial began with a small, white fixation cross centered in the middle of a black screen, presented for a randomly jittered amount of time between 750 ms and 1250 ms. This fixation cross would remain on the screen the entire trial, and participants were instructed to focus on it for the duration of the trial. Two flashing grey circles would then appear on the screen on either side of the fixation cross for a duration of 4000 ms. Participants were instructed to make a decision, denoted with a left or right shift key button press on a keyboard, about whether the right or left flashing circle looked brighter (or darker, depending on assigned counter-balanced condition). After a decision was made and the 4000 ms of flashing concluded (flashing would occur for the entire 4000 ms, regardless of a decision being made), there would be an inter trial interval of 500 ms, with only the fixation cross on the screen until the next trial would begin. Had a participant failed to make a choice during the 4000 ms of flashing, a fixation cross would remain on the screen until a choice was made, followed by the 500 ms inter trial interval. E2 involved an identical procedure to E1, except for minor adjustments to the timing and a limiting of frequencies tested to optimize the parameters for high-quality EEG data. One of these adjustments involved adding more time to both the initial fixation period and the inter trial interval (ITI), allowing for a time-frequency decomposition that included at least 500 ms before stimulus presentation and the entire duration of flickering (considering a wavelet convolution with 3 Hz as the slowest frequency). The amount of time the flashing circle was on the screen (4000 ms) remained constant between E1 and E2.

E1 consisted of 675 total trials per participant, broken into 27 blocks of 25 trials, with each block presenting all 25 conditions (see below) once in a random order, and E2 included 540 trials per participant, broken into 20 blocks of 27 trials, with each block presenting all 9 conditions (see below) 3 times in a pseudo-random order. Participants were instructed that they could self-time breaks between blocks, taking some brief time if they needed.

There was a total of 25 conditions in E1 consisting of different frequencies of the left and right flashing circles. The frequencies studied were 0 (e.g. constant / steady, or, no flashing), 4.4, 9.2, 13.3, and 17.1 Hz, with all frequency pairs presented, creating the 25 conditions. Note, since all pairs were presented, this means that each frequency, including the constant (0 Hz) stimuli, appeared equally often on both the left and right for every pair-type (e.g. both a 4.4 vs 9.2 Hz and a 9.2 vs 4.4 Hz pair appeared). The luminance value of all flickering circles was held constant, and remained grey on each flicker for 6 frames (50 ms) per flicker, irrespective of frequency. This meant a 4.4 Hz stimulus would flicker with 6 frames of grey followed by 21 frames of black (with 4 flickers of grey per second). As the number of frames per grey-circle flicker was held constant, only the number of frames of black differed to determine frequency, where 9.2, 13.3, and 17.1 Hz had 7 frames, 3 frames, and 1 frame of black, respectively, between 6 frames of grey. Participants were assigned to one of two conditions at the beginning of the experiment which informed whether they were asked to make a judgment about the brightness (n = 15) or darkness (n = 14) of the circles.

To allow for a number of trials sufficient for EEG analysis^40^, only 3 frequencies of stimuli were used in E2. These specific frequencies were chosen after reviewing the results of E1 and to specifically target frequencies within both the alpha and theta bands. The frequencies tested were 4.4, 9.2, and 13.3 Hz, with all frequency pairs presented, creating the 9 conditions. All other qualities about the stimuli were kept consistent with E1, including 2 experimental counterbalanced groups of brighter (n = 11) or darker judgments (n = 12). Also, as E2 incorporated EEG, extra time was taken to set up the cap on the participant, and clean up after the experiment. This made the experiment take approximately 2 hours, as opposed to 1.5 hours for E1.

### Acquisition

A BrainVision 32-channel active-wet electrode EEG was used to record EEG data sampled at 1000 Hz in a dark, quiet cement room. An online anti-aliasing filter was used of 280 Hz, and data were recorded DC coupled with no high pass filter. Data were digitized with 24 bits, with a bit depth of 0.0487 uV. Two electro-oculogram (EOG) measures were collected by aligning a pair of bipolar electrodes above and below the left eye (vertical EOG) and on the outside of each eye (horizontal EOG), grounded with their own ground electrode between the eyebrows. The data was referenced to an electrode on the right mastoid at the time of collection (online), and then re-referenced to an average of the left and right mastoid channels during data processing (offline). Scalp electrode impedances were held below 10 KΩ, and all 32 channels were recorded (10-20 system). EEG and EOG data were recorded using BrainVision Recorder software on a second computer running Windows 7. All datasets generated from both E1 and E2 are available from the corresponding author upon reasonable request.

### Pre-Processing

EEG data was processed using the EEGLAB toolbox^41^ in Matlab. Offline, data were low-pass filtered at 30 Hz, and high-pass filtered at 0.1 Hz. Epochs were removed locked to stimulus presentation, from 1250 ms before stimulus presentation and 4560 ms after stimulus presentation. Visual inspection of a participant’s epochs was used to look for any noisy channels involving abnormally large voltage changes or continuously noisy muscular artifact for a sustained amount of time (>10% of trials). Overall, 5 subjects’ data required the removal of one channel, and another 2 subjects’ data required the removal of two channels. No channels that were removed were part of the subset analysed. A manual epoch rejection was administered with the goal of removing any epochs that had unconventional artifacts, looking for anything that differed from a stereotypical blink, saccade or muscle artifact that could be easily removed with component analysis. An independent component analysis (runICA algorithm in EEGLAB) was then carried out on the epoched data, with a manual rejection of components that displayed stereotypical fronto-central artifact. An average of 13 components were removed for each individual. A second epoch rejection was manually completed after artifact components were removed – a final pass of the data to check the success of the ICA and to remove any artifact that still existed in the data after the first epoch rejection and component removal. All artifact rejection decisions were done by the first author after extensive training. After all data cleaning and processing, an average of 448 epochs (Range = 351-523) of an original 540 per participant, or 83%, remained and were usable for further analysis. Finally, all EEG data was spatially filtered with a surface Laplacian using the CSD toolbox^42,43^ in Matlab.

### Behavioural Data Acquisition and Processing

For both experiments, for each trial, behavioural data recorded in MATLAB consisted of a reaction time and a response side. We removed trials with reaction times less than 100 ms or greater than 3 standard deviations above each participant’s mean reaction time. For all participants in the darker condition, answers were inverted (i.e., a left choice became a right choice) so as to have all data in the form of the brighter condition for ease of analysis. The final, combined, EEG and behavioural data set aligned trials such that any trials removed during EEG were also removed from the trials for behavioural analysis and vice versa, and resulted in a final data set for each person with an average of 425 trials (Range = 294-521), or 79% (Range = 54-96%) of the original 540 trials.

### Statistical Procedure

Where we report the results of RMANOVAs we applied the Greenhouse Geisser correction for sphericity and report the corresponding epsilon value and corrected degrees of freedom. Main effects and interactions were considered significant if corrected p-values were less than 0.05. Significant two-factor interactions were followed up by running one-factor simple main effect RMANOVAs. Finally, significant simple main effect or main effect results were followed up by running all pairwise comparisons. These comparisons were considered significant if the Tukey-HSD corrected p-value was less than 0.05.

### EEG Measures and Analyses

All statistical EEG analyses were completed only on the occipital electrodes of O1 (left) and O2 (right). All analyses were carried out with scripts written and developed in Matlab including the EEGLAB^41^ toolbox.

#### ITC

Time frequency analyses involved transforming the original voltage data into frequency-specific information. We used the EEGLAB^41^ function newtimef() to perform a wavelet transform between frequencies of 3 to 30 Hz spaced out by .5 Hz, with a wavelet of 3 cycles for the lowest frequency (3 Hz), where cycles linearly increased with frequency by a rate of 0.5 cycles, reaching a 15 cycle wavelet at the highest frequency (30 Hz). Phase information in the form of ITC was considered, a measure between 0 and 1 of the regularity of a specific phase of a certain time and frequency, where, for example, when comparing ITC values between conditions, a higher ITC for one condition would be indicative of more event-related phase-locking in that subset of trials. This function provided ITC values for each subject, for each frequency-pair condition, for each electrode of interest (only O1/O2), for each frequency (3 to 30 Hz, by 0.5 Hz increments), and for each time point (with an estimate every 11.735 ms). For all ITC analyses, frequency bands of averaged ITC information were used, as opposed to a single frequency. These bands consisted of the closest frequency to the frequency of entrainment, and the closest frequency below and above it. For consistency and simplicity, we will refer to our ITC results as bands to acknowledge that the ITC information has been averaged over a range, or band, or frequencies. A 4.4 Hz band consists of an average of 4.0, 4.5, and 5.0 Hz ITC information, a 9.2 Hz band consists of an average of 8.5, 9.0, and 9.5 Hz ITC information, and a 13.3 Hz band consists of an average of 13.0, 13.5, and 14.0 Hz ITC information.

In general, for the first three results reported in the manuscript (see Figure 3) we averaged, tested, and reported ITC over a 1 s window from 1 – 2 s after stimulus onset. This window was chosen since it approximately straddles the mean response time for the group (1419 ms) where we hypothesized maximum neural biases would be evident. One exception to this window was made in calculating the “baseline” ITC levels which were taken between 500 and 200 ms before stimuli onset. In our fourth analysis, we binned our data by the behavioural response that was made. Since this creates a new grouping of trials, it was necessary to run a new time frequency decomposition to extract ITC values specific to each response condition. We now included only trials with same-frequency stimuli, and for each same-frequency pair, we extracted ITC values for the electrode contralateral to what was chosen as brighter and the electrode contralateral to what was chosen as dimmer. As reported, the key metric was the difference between the ITC contralateral-to-brighter and the ITC ipsilateral-to-brighter. Since we did not want to miss any potential temporally-evolving differences that were predictive of behaviour, this ITC was averaged over a longer window, stretching from 100 ms (fastest possible reaction time) to 3213 ms (closest time point from time frequency decomposition to two standard deviations above the mean reaction time on same-stimuli trials).

## Acknowledgements

The authors wish to thank M. Matta and A. Xue for their help in data collection. This work was supported by a Discovery Grant from the Natural Sciences and Engineering Research Council (RGPIN-2014-05248) to C. S. Chapman, and a graduate scholarship from the Alberta Gambling Research Institute to J. K. Bertrand.

## Author Contributions

J.B., K.M. and C.C. conceived of and designed these experiments. J.B. collected the data of these experiments. J.B. and C.C. wrote the main manuscript text and prepared all figures and the table. All authors (J.B., N.W., K.M., and C.C.) reviewed the manuscript.

## Additional Information

Competing financial interests: The authors declare no competing financial interests.

## REFERENCES

1. Von Helmholtz, H. Popular Lectures on Scientific Subjects (Longmans, Green, and Co., 1881).

2. Bartley, S. H. Subjective brightness in relation to flash rate and the light-dark ratio. J. Exp. Psychol. 23, 313–319 (1938).

3. Jensen, O., & Mazaheri, A. Shaping functional architecture by oscillatory alpha activity: Gating by inhibition. Front. Hum. Neurosci. 4, 1–8 (2010).

4. Palva, S., & Palva, J. M. Functional roles of alpha-band phase synchronization in local and large-scale cortical networks. Front. Psychol. 2, 1–15 (2011).

5. Klimesch, W., Sauseng, P., & Hanslmayr, S. EEG alpha oscillations: The inhibition–timing hypothesis. Brain Res. Rev. 53, 63–88 (2007).

6. Mathewson, K. E., Gratton, G., Fabiani, M., Beck, D. M., & Ro, T. To see or not to see: Prestimulus α phase predicts visual awareness. J. Neurosci. 29, 2725–2732 (2009).

7. Dugué, L., Marque, P., & VanRullen, R. The phase of ongoing oscillations mediates the causal relation between brain excitation and visual perception. J. Neurosci. 31, 11889–11893 (2011).

8. Mathewson, K. E., Fabiani, M., Gratton, G., Beck, D. M., & Lleras, A. Rescuing stimuli from invisibility: Inducing a momentary release from visual masking with pre-target entrainment. Cognition. 115, 186–191 (2010).

9. Mathewson, K. E. et al. Making waves in the stream of consciousness: Entraining oscillations in EEG alpha and fluctuations in visual awareness with rhythmic visual stimulation. J. Cog. Neuro. 24, 2321–2333 (2012).

10. De Graaf, T. A. et al. Alpha-band rhythms in visual task performance: Phase-locking by rhythmic sensory stimulation. PlOS One. 8, e60035 (2013).

11. VanRullen, R. Perceptual cycles. Trends Cogn. Sci. 20, 723–735 (2016).

12. Kohn, H., & Salisbury, I. Electroencephalographic indications of brightness enhancement. Vis. Res. 7, 461–468 (1967).

13. Dugué, L., Marque, P., & VanRullen, R. Theta oscillations modulate attentional search performance periodically. J. Cogn. Neurosci. 27, 945–958 (2015).

14. Cravo, A. M., Rohenkohl, G., Wyart, V., & Nobre, A. C. Temporal expectation enhances contrast sensitivity by phase entrainment of low-frequency oscillations in visual cortex. J. Neurosci. 33, 4002–4010 (2013).

15. Kawasaki, M., & Yamaguchi, Y. Effects of subjective preference of colors on attention-related occipital theta oscillations. NeuroImage. 59, 808–814 (2012).

16. Han, B., & VanRullen, R. Shape perception enhances perceived contrast: Evidence for excitatory predictive feedback? Sci. Rep. 6, 22944; doi:10.1038/srep22944 (2016).

17. Han, B., & VanRullen, R. The rhythms of predictive coding? Pre-stimulus phase modulates the influence of shape perception on luminance judgments. Sci. Rep. 7, 43573; doi:10.1038/srep43573 (2017).

18. Zhu, W., Drewes, J., & Melcher, D. Time for awareness: The influence of temporal properties of the mask on continuous flash suppression effectiveness. PLOS One, 11, e0159206; doi:10.13171/journal.pone.0159206 (2016).

19. Jahn, T. L. Brightness enhancement in flickering light. Psychol. Rev. 51, 76–84 (1944).

20. Kelly, D. H. Nonlinear visual responses to flickering sinusoidal gratings. J. Opt. Soc. Am. 71, 1051–1055 (1981).

21. Bartley, S. H., Paczewitz, G., & Valsi, E. Brightness enhancement and the stimulus cycle. J. Psychol. 43, 187–192 (1957).

22. van der Horst, G. J., & Muis, W. Hue shift and brightness enhancement of flickering light. Vis. Res. 9, 953–963 (1969).

23. Pantle, A. Flicker adaptation—II. Effect on the apparent brightness of intermittent lights. Vis. Res. 12, 705–715 (1972).

24. Magnussen, S., Bjørklund, R. A., & Krüger, J. The perception of flicker: Theoretical note on the relation between brightness and darkness enhancement. Scand. J. Psychol. 20, 257–258 (1979).

25. Kurtenbach, A., Rüttiger, L., Kaiser, P. K., & Zrenner, E. Influence of luminance flicker and purity on heterochromatic brightness matching and hue discrimination: A postreceptoral opponent process. Vis. Res. 37, 721–728 (1997).

26. Palva, S., & Palva, J. M. Functional roles of alpha-band phase synchronization in local and large-scale cortical networks. Front. Psychol. 2, 204 (2011).

27. Klimesch, W. Alpha-band oscillations, attention, and controlled access to stored information. Trends Cogn. Sci. 16, 606–617 (2012).

28. Klimesch, W. EEG alpha and theta oscillations reflect cognitive and memory performance: A review and analysis. Brain Res. Rev. 29, 169–195 (1999).

29. Van Dijk, H., Schoffelen, J. M., Oostenveld, R., & Jensen, O. Prestimulus oscillatory activity in the alpha band predicts visual discrimination ability. J. Neurosci. 28, 1816–1823 (2008).

30. Agarwal, G. et al. Spatially distributed local fields in the hippocampus encode rat position. Science. 344, 626–630 (2014).

31. Mathewson, K. E. et al. Pulsed out of awareness: EEG alpha oscillations represent a pulsed-inhibition of ongoing cortical processing. Front. Psychol. 2, 1–15 (2011).

32. Benedetto, A., Spinelli, D., & Morrone, M. C. Rhythmic modulation of visual contrast discrimination triggered by action. Proc. R. Soc. B. 283, 20160692; doi:10.1098/rspb.2016.0692 (2016).

33. Tomassini, A., Spinelli, D., Jacono, M., Sandini, G., & Morrone, M. C. Rhythmic oscillations of visual contrast sensitivity synchronized with action. J. Neurosci. 35, 7019–7029 (2015).

34. Hanslmayr, S. et al. Visual discrimination performance is related to decreased alpha amplitude but increased phase locking. Neurosci. Lett. 375, 64–68 (2005).

35. Holcombe, A. O., & Chen, W. Y. Splitting attention reduces temporal resolution from 7 Hz for tracking one object to < 3 Hz when tracking three. J. Vis. 13, 1–19 (2013).

36. Macdonald, J. S., Cavanagh, P., & VanRullen, R. Attentional sampling of multiple wagon wheels. Atten. Percept. Psychophys. 76, 64–72 (2014).

37. Crouzet, S. M., & VanRullen, R. The rhythm of attentional stimulus selection during visual competition. Preprint at http://dx.doi.org/10.1101/105239 (2017).

38. Chaumillon, R., Blouin, J., & Guillaume, A. Interhemispheric transfer time asymmetry of visual information depends on eye dominance: an electrophysiological study. Front. Neurosci. 12, 1–19 (2018).

39. Brainard, D.H., & Vision, S. The psychophysics toolbox. SpatialVision. 10, 433–436 (1997).

40. Cohen, M.X. Analyzing Neural Time Series Data: Theory and Practice (MIT Press, 2014).

41. Delorme, A., Makeig, S. EEGLAB: an open source toolbox for analysis of single-trial EEG dynamics. J. Neurosci. Methods. 134, 9–21 (2004).

42. Kayser, J., & Tenke, C.E. Principal components analysis of Laplacian waveforms as a generic method for identifying ERP generator patterns: I. Evaluation with auditory oddball tasks. Clin. Neurophysiol. 117, 348–368 (2006).

43. Kayser, J., & Tenke, C.E. Principal components analysis of Laplacian waveforms as a generic method for identifying ERP generator patterns: II. Adequacy of low-density estimates. Clin. Neurophysiol. 117, 369–380 (2006).

